# Multiband acceleration can provide moderate improvements in single-subject voxel-wise statistics in block-design task-based fMRI

**DOI:** 10.1101/756361

**Authors:** Ritu Bhandari, Valeria Gazzola, Christian Keysers

## Abstract

Multiband (MB) acceleration of functional magnetic resonance imaging has become more widely available to neuroscientists. Here we compare MB factors of 1, 2 and 4 while participants view complex hand actions vs. simpler hand movements to localize the action observation network. While in a previous study, we show that MB4 shows moderate improvements in the group-level statistics, here we explore the impact it has on single subject statistics. We find that MB4 provides an increase in *p* values at the first level that is of medium effect size compared to MB1, providing moderate evidence across a number of voxels that MB4 indeed improves single subject statistics. This effect was localized mostly within regions that belong to the action observation network. In parallel, we find that Cohen’s *d* at the single subject level actually decreases using MB4 compared to MB1. Intriguingly, we find that subsampling MB4 sequences, by only considering every fourth acquired volume, also leads to increased Cohen’s *d* values, suggesting that the FAST algorithm we used to correct for temporal auto-correlation may over-penalize sequences with higher temporal autocorrelation, thereby underestimating the potential gains in single subject statistics offered by MB acceleration, and alternative methods should be explored. In summary, considering the moderate gains in statistical values observed both at the group level in our previous study and at the single subject level in this study, we believe that MB technology is now ripe for neuroscientists to start using MB4 acceleration for their studies, be it to accurately map activity in single subjects of interest (e.g. for presurgical planning or to explore rare patients) or for the purpose of group studies.

## 1. Introduction

Functional magnetic resonance imaging (fMRI) is increasingly used as a non-invasive pre-surgical tool for planning function-preserving neurosurgeries for epilepsy and brain tumor patients (Lindquist & Mejia, 2015; Nadkarni et al., 2014; Ni et al., 2019; Wengenroth et al., 2011). Given the extreme importance of the outcome from such examinations, it is critical to use a scanning sequence that would result in the most robust estimations of activation in different brain regions. Multiband (MB) accelerated scanning that allows the acquisition of magnetic resonance imaging (MRI) signals from more than one spatial coordinates at a time has been tested in over 40 studies with resting state and task-based fMRI (for a detailed overview see Bhandari et al., 2020). The advantages offered by this acquisition scheme that affords higher sampling rate have been shown in many resting state studies (Feinberg et al., 2010; Griffanti et al., 2014; Koopmans et al., 2012; Preibisch et al., 2015). Several task based fMRI studies also show comparable or better results for MB accelerated sequences (Chen et al., 2015; Demetriou et al., 2018; Moeller et al., 2010; Sahib et al., 2018; Todd et al., 2017). In our recent study we analysed task based fMRI data acquired with multiple sequences with different MB+SENSE acceleration factors and showed that sequences with MB acceleration perform better in terms of voxel-wise group-level *t*-values (Bhandari et al., 2020). This was one of only two studies (Boyacioĝlu et al., 2015) that quantified and compared the effects of MB acceleration on a *group level*, voxel-wise statistics. Most other studies that assessed task correlated BOLD signals for quantifying performance of MB acceleration, concluded their findings using single subject level statistics, extracting *t*-values within regions of interest (ROI) (Kiss et al., 2018; McDowell & Carmichael, 2018; Sahib et al., 2016, 2018; Todd et al., 2016). Here we aim to contribute to that effort to characterize the potential benefits of MB acceleration to voxel-wise signals at the individual level, by analysing the effect of MB not only in specific ROIs that were activated by a task, but throughout the entire brain. We therefore computed within-subject ANOVAs that directly contrast statistical values between sequences to reveal if there is a consistent improvement in the ability to detect a task network with increased acceleration.

An important consideration to include while directly comparing the subject level statistics coming from different MB acceleration sequences is that if one keeps acquisition time constant, higher MB will generate more numerous samples. The subject-level *t*-values and the critical *t*-values corresponding to a particular *p*-value dependent on the number of samples. Therefore, what value should be compared across different MB sequences merits careful consideration.

The parameter estimate (or β maps) are not affected by the sampling rate. However, these cannot be used while comparing MB acceleration as one of the side effects of accelerated sampling is the loss in grey-white contrast in the functional images (Bhandari et al., 2020). This paired to the default global normalization in SPM, which is necessary to homogenize the data coming from different subjects and groups, results in a spurious difference across acquisition schemes due to different grey-white contrast that result from the shorter acquisition time, which in turn leads to less T1 recovery.

Intuitively, given that one is usually ultimately thresholding fMRI maps using a specific *p*-value, an obvious measure on which to compare different sequences would be the voxel-wise *p*-value. Because *p-* values are not normally distributed, to include them in an ANOVA, one would then *z*-transform the *p-* values. Given the known degrees of freedom and the *t*-value per voxel, one can calculate *p*-values and then convert them to *z*-values to make the results normally distributed and allow direct comparison of data between sequences within the same statistical model.

Alternatively, one can look at the “effect-size” *d*-values, which requires the normalization of the *t*-values by dividing them by the square root of the degrees of freedom. The *d* transformation normalizes the *t*-values for the difference in the degrees of freedom, thereby also allows direct comparison.

Figure 1 visualizes the impact of sample size on *t*, *p, z* and *d* values. For this aim, we drew a vector of 1000 numbers from a μ=0.1 and σ=1 random normal distribution to simulate an effect size of δ=0.1, and then performed one-sample *t*-tests against zero considering the first *n* elements of that vector. As can be seen, *t*, *p* and *z* values increase along slightly differently shaped curvilinear gradients with increasing *n*, while *d* remains constant, close to δ. This is because *t, p* and *z* values are all metrics of the evidence we have for an effect, and evidence increases with sample size, while *d* is a metric of the size of the effect, which in our simulation is constant. While this simulation confirms that increasing the number of samples should lead to better *t* and *p* values, it does not take into account (a) the increased noise due to the inefficient separation of the simultaneously acquired voxels with MB acquisition and (b) the increased temporal auto-correlation with higher sampling rate. We therefore perform direct comparison of *z* and *d* values associated with different MB sequences to see if the advantages of higher sampling surpass the negative effects of the increased noise and noise correlation.

**Figure 1:**
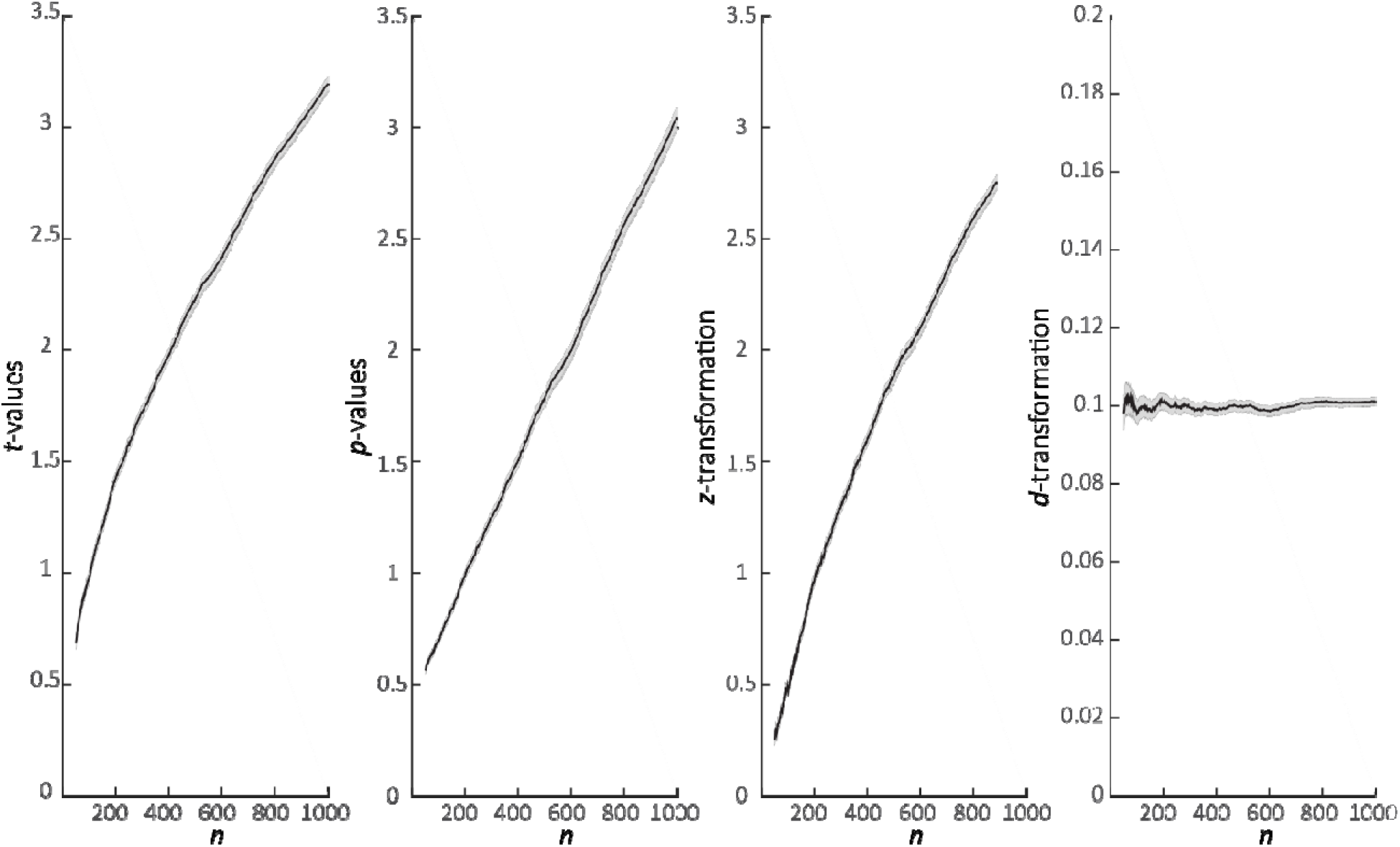
Effect of sample size on *p, t, z* and *d* values. We drew a vector of 1000 numbers from a μ=0.1 and σ=l random normal distribution to simulate an effect size of δ=0.1, and calculated the *p, t, z* and *d* value of a *t*-test comparing the first *n* of these numbers against zero, varying *n* from 50 to 1000. The *z* values were calculated by performing the normal inverse of the Student’s *t* cumulative distribution function. The *d* values were calculated as *t*/√(*n*-1). The graphs plot the mean±sem (over the 1000 simulations) of these values as a function of the number of samples (*n*) considered. For *p*, the values are plotted on a −log10 scale. Note how for a given effect size (cohen δ=0.1) increasing sample size increases *p, t, z* values, while *d* values remain stable at a value equal to the simulated effect size.

## 2. Methods

### Subjects, Task and Acquisition

Data were acquired from 24 healthy volunteers (two were excluded due to missing data or technical errors, *N* =22), on a 3T Philips scanner, using a commercial version of Philips’ MB implementation (based on software release version R5.4). A 32-channel head coil was used. Functional data were acquired using different acquisition sequences (see table 1). Two types of stimuli were used: Complex actions (CA) showed the hand interacting with the object in typical, goal directed actions. For example, the hand of the actor reached for the lighter placed on the table, grasped it, and lit the candle with it. Complex controls (CC) stimuli had the exact same setting as the CA but the actor’s hand did not interact with or manipulate the object on the table, instead, made aimless hand movements. A block was composed of three movies of the same category (CA or CC) and lasted 7s. Each fMRI session was composed of 13 blocks per stimulus category for a total of 26 blocks, presented in a randomized order. The inter-block-interval lasted between 8 – 12s and consisted of a fixation cross on a gray and blue background similar to the stimuli background. These sessions were presented to each subject multiple times, showing the same blocks but in different order, with each session acquired with a different acquisition sequence, varying in MB factor (table 1). The order of acquisition was randomized between subjects. Importantly, the duration of the sessions was similar (~8 min), but more functional volumes were acquired during sessions at lower TRs. Two additional sequences were acquired (a 3×3×3.3mm, and a 2.7mm isotropic sequence with MB4 but SENSE 1.5). The former is used here to generate a Action Observation Network (AON) localizer (see below), but is not used in the comparison of MB, because this sequence differs from the other sequences in terms of voxel size. The latter is not used here, because its sense value, 1.5, differs from that of the other values, 2. (for details about those sequences see Bhandari et al., 2020).

**Table 1.** Parameters used for scanning.

| Parameters | TR | 2.45s | 1.22s | 0.63s |
| --- | --- | --- | --- | --- |
| Multiband factor (MB) |  | 1 | 2 | 4 |
| SENSE acceleration factor (S) |  | 2 | 2 | 2 |
| Acquired voxel size (mm) |  | 2.7 isotropic | 2.7 isotropic | 2.7 isotropic |
| Flip angle in degrees |  | 79 | 64 | 50 |
| Number of slices |  | 44 | 44 | 44 |
| Acquired volumes |  | 200 | 400 | 780 |
| Slice gap (mm) |  | 0.27 | 0.27 | 0.27 |
| Field of view (mm) |  | 216x216x130.4 | 216x216x130.4 | 216x216x130.4 |

### Preprocessing

Data were preprocessed using SPM12 (Wellcome Trust Centre for Neuroimaging, UCL, UK) with Matlab version 8.4 (The MathWorks Inc., Natick, USA). Briefly, functional images were slicetime corrected and then realigned to the estimated average. Anatomical images were co-registered to the mean functional image, and segmented. The normalization parameters that were generated during segmentation were used to bring all the images to the MNI space. The resampled voxel size for the functional images was 2 × 2 × 2 mm and 1 x 1 x 1 for the anatomical scans.

### Subject-level GLM

Subject level GLM included CA and CC as two separate task predictors with each predictor having 13 blocks of 7s. Regressors included the six motion parameters estimated during realignment, first five principal components of cerebrospinal fluid and five principal components of white matter (total 16 regressors). As by default, the GLM involves a global normalization step, described in the SPM manual (https://www.fil.ion.ucl.ac.uk/spm/doc/spm12_manual.pdf), section 8.7 as “SPM computes the grand mean value 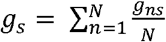. This is the fMRI signal averaged over all voxels within the brain and all time points within session s. SPM then implements ‘Session-specific grand mean scaling’ by multiplying each fMRI data point in session s by 100/g_s_.”. The model included a low-pass filter at 128s and autoregressive correction was performed using the FAST option. We chose to use the FAST option as previous studies have suggested that FAST is better than AR(1) at correcting statistics for the autocorrelation in fast data acquisition schemes where autocorrelation extends beyond an autoregressive order of 1 because of the short TR (Bollmann et al., 2018; Todd et al., 2016). The *t*-contrast of interest was always CA-CC which activates the Action observation network (AON, Gazzola & Keysers, 2009, Figure 3B). Only in Figure 5, we used AR(1).

To explore the effect of the sampling, we performed two additional subject-level GLMs, one for sequences with MB factor 2 and one for MB factor 4, where we decimated the data to include only the odd data points for MB2 and every 4^th^ data point in MB4 sequence. This resulted in inclusion of 200 data points for MB2 and 195 data points for MB4 sequence. In the SPM model design, the TR was changed to 1.22*2 and 0.63*4 and the microtime resolution was changed from the default of 16 to 16*2 and 16*4 for MB2 and MB4 sequences respectively. All the other settings were as described above. Additionally, we re-an the analysis with all the above settings, but instead of the FAST we used the AR(1) for correcting the autocorrelation.

### z-transformation

For each subject, the *z*-transformation was applied to the SPM *t*-maps of the CA-CC contrast. We read out the degrees of freedom estimated by SPM while visualizing the CA-CC contrast. SPM *t*-maps were then converted into *z*-maps using the normal inversion of the student’s *t* – cumulative density in matlab using the the equation *z=norminv*(*tcdf*(*t, df*), *μ σj*/ The assigned values for μ an σ were 0 and 1 respectively. The *z*-values for higher *t*-value (~ *t* = 5.2) were estimated as infinite using the above function. These *z*-values were assigned to be 9.

### d-transformation

For the *d*-transformation, we read out the degrees of freedom estimated by SPM while visualizing the CA-CC contrast, and then using the formula *d=t*/√(*df*), the *t*-maps were converted to *d*-maps.

### Group-level GLM

To identify the effect of MB acceleration on voxel-wise, subject-level statistics, two within-subject group-level ANOVAs were performed, one for *z* and one for *d* values. These group level ANOVAs assess whether in any voxels, the *z* (or *d*) value at the single subject level systematically changes as a function of MB. Six pairwise contrasts were computed between the three sequences (MB1>MB2, MB1>MB4, MB2>MB4, MB2>MB1, MB4>MB1, MB4>MB2). All the results were evaluated at *q_fdr_*<0.05 and *p_uncorr_*<0.001 and cluster threshold of 50 voxels.

Two additional within-subject ANOVAs for the full vs. decimated dataset were also computed separately for MB2 and MB4 sequences. Full>decimated and decimated>full contrasts were computed. All the results were evaluated at q_fdr_<0.05 and cluster threshold of 50 voxels.

### AON Localizer

To address whether gains in *z* or *d* values as a function of MB fall within regions activated by our task or not, we created a AON mask (18360 voxels) using the results of a group-level analysis for the contrast CA-CC of the same subjects but from a sequence with no MB and bigger voxel size (3×3×3.3mm). We used the same first level GLM as described above, but then brought the contrast CA-CC of each participant to a second level *t*-test, and thresholded results at *q_fdr_*<0.05 and cluster threshold of 50 voxels. The mask is shown as gray outlines in Figures 2, 4 and 5, and in blue in Figure 3.

**Figure 2:**
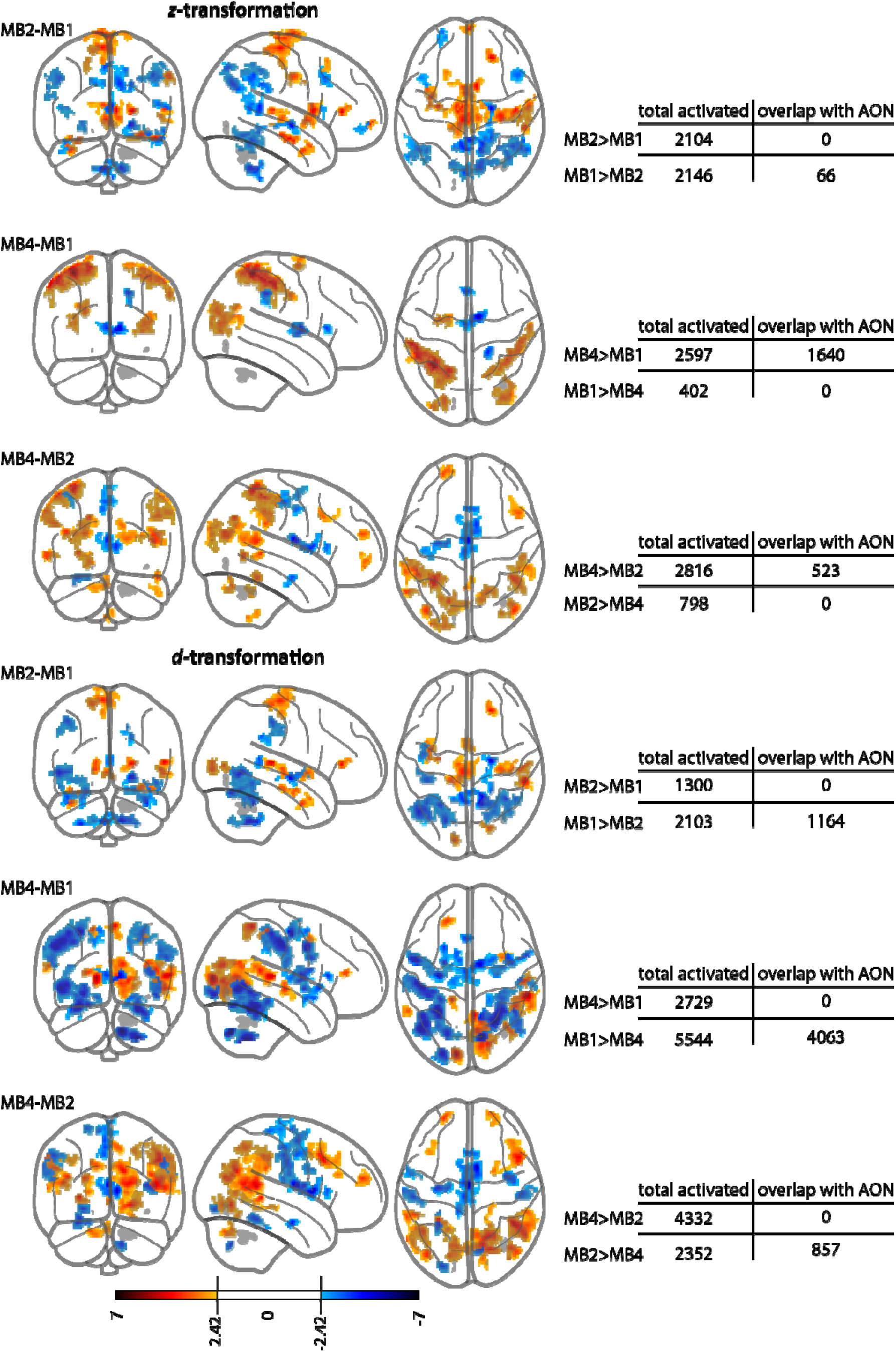
Pairwise contrasts between the three sequences in terms of *z* and *d* values. Row 1-3 for *z* and 4-6 for *d* values. Hot colours represent voxels where faster is better (i.e. higher MB>lower MB) and the cold colours are the opposite contrast. All the contrasts are evaluated at *p_uncorr_*<0.001 and a cluster threshold of 50 voxels. In grey are the areas that belong to the CA-CC network. It is derived from the group analysis of the same subjects but the data was acquired using a standard sequence with no MB and bigger voxel size (3×3×3.3mm) at *q_fdr_*<0.05.

**Figure 3:**
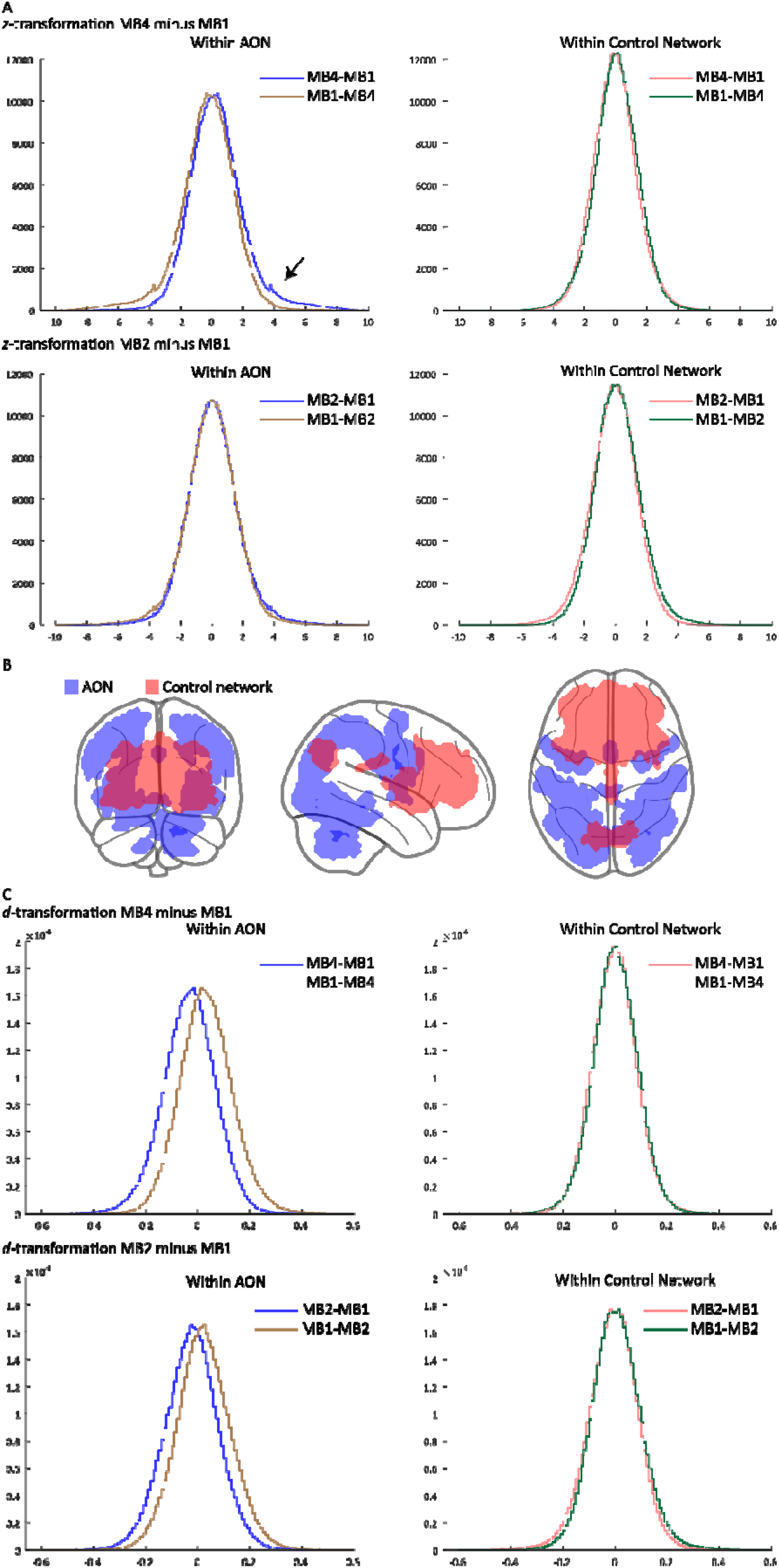
Voxelwise effect of MB on *z* and *d*-values. (a) Histogram within AON (left) and within a control network (right) showing MB4/MB2-MB1 contrasts for *z*-transformation. The arrow in the top pannel points to the slightly higher *z*-values For MB4-MB1 contrast within the AON but not in the control network, (b) Blue shows the task network (18360 voxels) and in orange is the control network of similar size (18309 voxels). The overlap between these two networks is minimal (70 voxels). (c) Histogram similar to (figure 3a) but now for *d*-values. Both for MB4-MB1 and MB2-MB1, the histogram reveals that MB1 performs better in the AON (brown histogram shifted to positive side compared to the blue one) but not in control network.

As a control network, we used a resting state network of a similar size (18309 voxels) which was also derived from the same subjects using the sequence with no MB and bigger voxel size (3×3×3.3mm). Briefly, the component (component 8 as described in Bhandari et al., 2020) was identified for each subject using their residual time series (after removig task-correlated BOLD) and performing a dual regression with the resting state masks presented in Smith et al., 2009. The individuals’ component 8 maps were then brought to a group level analysis and threshholded at *p_FWE_*<0.05 and cluster threshhold of 50 voxels. This resulted in a mask similar in size to the AON mask with minimum overlap (70 voxels) with the AON mask. This mask is shown in orange in Figure 3.

## 3. Results

For the *z* values, the pairwise contrasts between MBs showed no significant results at *q_FDR_*<0.05. However, lowering the threshold to *p_uncorr_*<0.001 showed that in regions that show increases of activity in the CA-CC contrast at the group level (i.e. within the Action Observation Network, AON) MB4 showed higher *z* values compared to MB1 and MB2 (see Figure 2, rows 1-3). There were 1640 voxels (at *P_uncorr_*<0.001) for the MB4-MB1 and 523 (at *p_uncorr_*<0.001) voxels for MB4-MB2 contrasts within the AON with higher *z*-values. The opposite contrast (i.e. MB1-MB4 and MB2-MB4) showed no significant voxels within the AON. This shows that while the benefits of MB on the significance of subject-level voxel-wise analysis are too modest to survive correcting for multiple comparisons across the entire brain, at a lower threshold, sequences with MB4 do increase *z* values (and hence, *p* values) in regions that are task-activated at the group level. The difference between MB1 and MB2 was not as well defined and within the AON, MB1 performed slightly better with 66 voxels showing better *z*-values for MB1-MB2, whereas the MB2-MB1 contrast did not show any clusters in the AON.

For the *d* transformation, the contrast MB1-MB4 showed five clusters (452 voxels, at *q_fdr_*<0.05) within the AON that were significant at *q_fdr_*<0.05 (Table 2) suggesting that the sequences *without* MB show higher effect sizes than sequences with MB. There were no other significant clusters outside the AON. Lowering the threshold to *p_uncorr_*<0.001 showed a similar trend: lower MBs performed better than the higher ones with more significant voxels within the AON (see Figure 2, rows 4-6). While MB2>MB1, MB4>MB1 and MB4>MB2 showed no significant clusters within the AON, MB1>MB2, MB1>MB4 and MB2>MB4 showed 1164, 4063 and 857 voxels respectively within the AON.

**Table 2.**
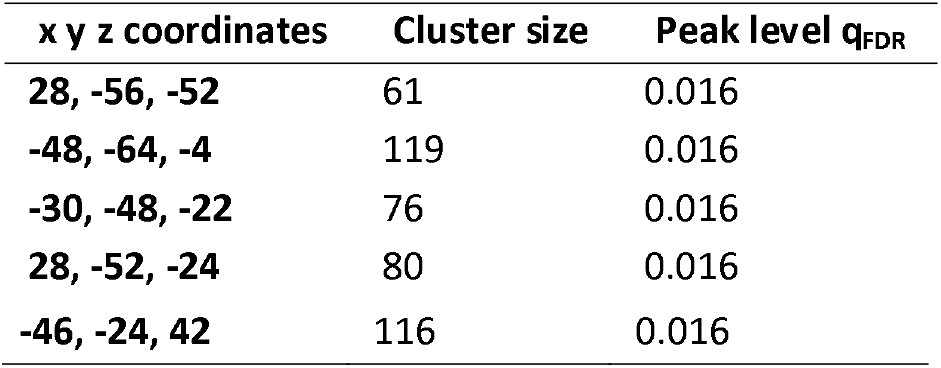
Coordinates of the five clusters that survive *q_FDR_*<0.05 for the contrast MB1-MB4 comparing *d*-values.

To further illustrate the effect of MB on *z* values within or outside of the AON, for each voxel of each participant, we subtracted the *z* value MB4-MB1, and plotted the histogram of these differences across participants and voxels within and outside the AON network. We then did the same for MB2-MB1. Figure 3A left shows these results in blue within the AON and on the right it shows these results in orange within the control network. To better visualize whether these histograms are shifted towards the positive side, we also show the histogram for MB1-MB4 or MB1-MB2 in brown (for AON) and green (for control network). The shift we see, with the blue histogram showing more highly positive values than the brown (pointed by the arrow in figure 3A) illustrates the systematic tendency towards higher *z* values in MB4 than MB1 within the AON. Replicating the same analysis in a control network of a similar size (a resting state network) does not show the same effect, illustrating that the gains obtained using MB4 are localized in regions associated with the task. This provides further insight into the source of improvement seen at group level statics with MB (as reported in Bhandari et al., 2020) showing that while the benefit of MB acceleration is small (only visible at an uncorrected level) and non-significant (at *q_FDR_*<0.05) when considering a voxel at a time, it may add up to show benefits at larger scales.

We repeated this with the *d*-values confirming the results in figure 2 that for the *d*-values, sequences without MB show higher values than MB4 and MB2 within the AON, but not in a network of a similar size (Figure 3C).

Finally, when comparing sequences with and without MB, two factors play a role. MB increases the number of samples but also alters the properties of the images. Increasing MB for instance increases noise in the images (reduced raw tSNR) and reduced grey white matter contrasts (CNR) (Bhandari et al., 2020). Increased noise could explain why MB decreases *d*, which can be conceived of as a metric of how much information can be drawn per image. More samples could explain the increase in *z*, which is a metric of the evidence for an effect drawn from the entire set of images acquired. To isolate the effect of the increased number of samples, we also compared sequences within a given MB sequences in which we reduced the number of samples by decimating the sequences, i.e. keeping every other volume at MB2 or keeping only every forth sample at MB4. Since all the other image aspects are identical in these full vs decimated data, we expected the full data to show higher *z* but identical *d* values compared to the decimated sequence. For the *z* values we however found no significant differences between the full and decimated versions of MB2 and MB4 data at *q_FDR_*<0.05. Decreasing the threshold to *p_uncorr_*<0.001 showed that MB4 full dataset indeed performs better than MB4 decimated dataset, as expected (Figure 4, second panel). Surprisingly, for the *d* values, for both MB2 and MB4 sequences, the decimated data performed significantly better than the full data at *q_FDR_*<0.05 and *p_uncorr_*<0.001 (Figure 4, panels 3-6).

**Figure 4:**
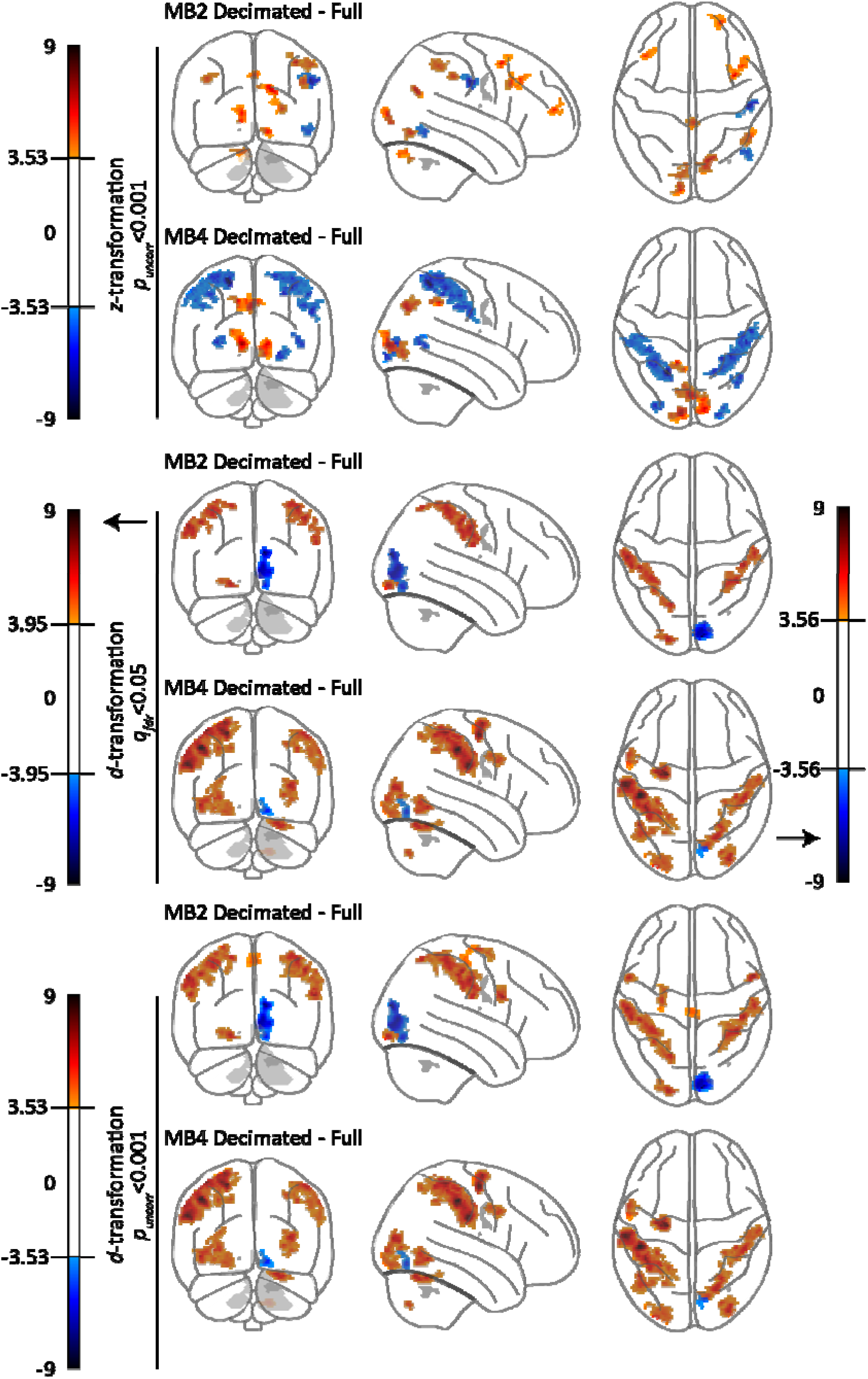
Effect of decimation on *z* and *d*-values. Top two panels present pairwise contrasts of *z* values for decimated – full data of MB2 and MB4 respectively. These contrasts are evaluated at *p_uncorr_*<0.001, a cluster threshold of 50 voxels. Middle two panels present these contrasts for d-values at *q_fdr_*<0.05, a cluster threshold of 50 voxels and lower two pannels present these contrasts for *d*-values at *p_uncorr_*<0.001, a cluster threshold of 50 voxels. Warm colors indicate voxels where the decimated sequence have higher *d* values than the full sequence, cold colours show the opposite. In grey are the areas that belong to the AON. It is derived from the group analysis of the same subjects but the data was acquires using a standard sequence with no MB and bigger voxel size (3×3×3.3mm) at *q_fdr_*<0.05.

## 4. Discussion

The aim of this study was to investigate whether the voxel-wise statistics of single subjects improve from using MB accelerations. Twenty two participants took part in a block-design study identifying the Action Observation network (AON) by contrasting blocks showing movies of complex goal-directed actions (CA) against blocks showing movies of complex control (CC) movements of the hand not directed at actions. Each participant performed this experiment while brain activity was measured using MB1, MB2 and MB4. For each participant and voxel, we then performed a *t*-test assessing the contrast of interest (CA-CC). We then explored whether higher MB lead to more significant *t*-tests across our participants.

As mentioned in the introduction, comparing the parameter estimates across MB is not meaningful, due to the differences in gray-white-matter contrast across the different MB (due to the differences in TR), and comparing *t*-values is also problematic. Accordingly, because scientists typically assess single subject results by thresholding at a specific *p* value, we first transformed the *t*-values obtained at each MB into their *p*-values and then *z*-transformed these to have a normally distributed proxy of significance. We found that in particular within regions that are activated by our task of interest (i.e. within the AON), we see that MB4 leads to moderately higher *z* and hence *p* values compared to MB1. MB2 only generates very modest improvements compared to MB1. These increases are not strong enough to survive a whole brain FDR correction, but are apparent in and restricted to voxels with task relevance (i.e. within the AON), when reducing the threshold to *t*=2.42, corresponding to a medium effect size of *d*=0.45, which generates at least moderate evidence for an improvement using a Bayesian analysis (*BF*_+0_=4.6, usingjasp-stats.org summary statistics function). If available, we can thus recommend the use of MB4 to perform task-based fMRI analysis at the single subject level. This confirms previous studies that used summary statistics with task based fMRI to show that the results of MB acceleration are comparable or slightly better (Boubela et al., 2014; Boyacioĝlu et al., 2015; Moeller et al., 2010).

We also calculated the effect size *d* of the contrast CA-CC as another way to comparing the data from sequences with different sample sizes. This was done by dividing the *t*-values with the square root of the degrees of freedom. We found, that *d* decreases with increasing MB, in line with the reduction in raw tSNR found in our data (Bhandari et al., 2020). That MB4 increases *z* values despite reducing *d* values shows that the increase in sample size suffices to counteract the slightly reduced image quality assessed by *d.*

To further explore the trade-off between sample size and image quality, we decided to decimate MB4 sequences, keeping only every fourth sample, and compare the *z* and *d* values with those of the full MB4 sequence. That way, the quality of the images considered in the analysis is constant but their quantity differs. We thus expected *d* to be the same, but *z* to be increased in the full compared to the decimated sequence. Intriguingly, this was not what we found. For *z*-values, full-decimated contrast showed no significant voxels at *q_fdr_*<0.05, but did show significant voxeles at *p_uncorr_*<0.001 within th AON for MB4. Moreover, *d* values were higher for the decimated sequences than for the full sequence (within the AON), mirroring the effect we had found between MB1 and MB4. It is unclear to us why this phenomenon arises. In its calculation of the *t*-value, SPM penalizes results for the serial autocorrelation of the data. This autocorrelation will be higher for samples taken closer together. By decimating the data, we reduced the autocorrelation of the data. If this penalization were optimal, dividing the *t*-values by the effective *df* calculated by SPM should have lead to similar *d* values for the full and decimated sequence (see Figure 1), but instead, *d* values decrease in the full sequence suggesting that SPM overpenalizes the *t*-statistics in our instance. This raises the intriguing possibility, that the advantages of MB might actually be underestimated by the proceedures currently performed in SPM using the FAST method for correction for serial autocorrelation. To test if this reasoning could be true, we performed the analysis of the full vs decimated sampling with the AR(1) instead of the FAST autocorrelation modelling implemented in SPM12. The results confirmed our intuition that FAST may be over-conservative. We found that for *z*-transformation, the full – decimated contrast shows higher values within the AON at *q_FDR_*<0.05 (figure 5) and for (*d*-transformation the full-decimated contrast showed no significant clusters at *q_fdr_*<0.05 suggesting comparable results and lowering the threshold to *p_uncorr_*<0.001 only showed a small 59 voxel cluster for decimated – full contrast at MNI co-ordinates −22 – 12 60. In light of these findings, we invite the community to explore subsampling as a method to benchmark the ability of analysis packages to deal with autocorrelation.

**Figure 5:**
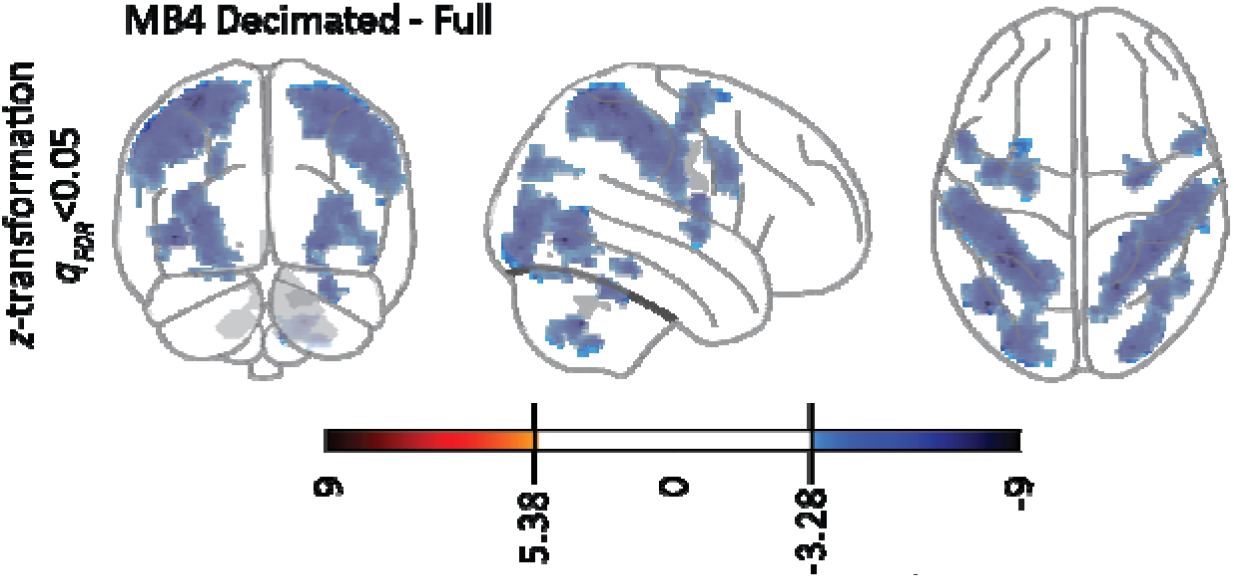
Impact of AR(1) on the results of decimated data. Pairwise contrasts of *z* values for decimated – full data of MB4 sequence. Warm colors would indicate voxels where the decimated sequence has higher *z* values than the full sequence, cold colours show the opposite. Contrast are evaluated at *q_fdr_*<0.05 and a cluster threshold of 50 voxels. In grey are the areas that belong to the AON.

We would like to mention a number of limitations of our study. First, our *z* transformation is slightly hampered by the fact that for high *t* values (>5.2), *p* values become so small, that the *z*-transformation generates infinite results in matlab. We then set these results to *z*=9, which could slightly bias our results. Second, while we compared the AR(1) and FAST autocorrelation algorithms implemented in SPM12, we did not systematically compare other software packages for correcting for serial autocorrelation. However, given our findings we believe that exploring alternative methods of dealing with temporal autocorrelation would be important to optimize analysis pipelines and reap the advantages of MB. Thirdly, we used compcorr regressors for de-noising the data, but did not assess alternative ways of cleaning such as FIX, which may provide additional benefits for accelerated sequences.

In summary, however, we find that using MB4 provides moderate improvements of the *p*-values that can be obtained in single subjects, and feel that our use of a comparatively larger sample size and subtle contrast between two high-level visual stimuli may provide information that is of value for the community of cognitive neuroscientists interested in the use of MB. Together with our evidence that MB4 provides significant benefits for group level analysis (Bhandari et al., 2020), we therefore believe that MB technology has reached a level where cognitive neuroscientists can benefit from its use.

## Acknowledgements

This work was supported by the Netherlands Organization for Scientific Research (VIDI: 452-14-015 to V.G.), the Brain and Behavior Research Foundation (NARSAD young investigator 22453 to V.G.), the European Research Council of the European Commission (ERC-StG-312511 to C.K.) and the BIAL foundation grant (255/16 to V.G., C.K. & R.B.). We thank Spinoza centre for neuroimaging where the scanning was performed and the staff members of Spinoza centre.

## Conflict of Interest

The authors report that there is no conflict of interest.

## Notes

### Competing Interest Statement

The authors have declared no competing interest.

